# Isochoric supercooled preservation and revival of human cardiac microtissues

**DOI:** 10.1101/2021.03.22.436466

**Authors:** Matthew J. Powell-Palm, Verena Charwat, Berenice Charrez, Brian Siemons, Kevin E. Healy, Boris Rubinsky

## Abstract

Low-temperature *ex vivo* preservation and tissue engineering based on human induced pluripotent stem cells (hiPSC) represent two of the most promising routes towards on-demand access to organs for transplantation. While these fields are often considered divergent from one another, advances in both fields present critical new opportunities for crossover. Herein we demonstrate the first-ever sub-zero centigrade preservation and revival of autonomously beating three-dimensional hiPSC-derived cardiac microtissues^1^ via isochoric supercooling^2^, without the use of chemical cryoprotectants. We show that these tissues can cease autonomous beating during preservation and resume it after warming, that the supercooling process does not affect sarcomere structural integrity, and that the tissues maintain responsiveness to drug exposure following revival. Our work suggests both that functional three dimensional (3D) engineered tissues may provide an excellent high-content, low-risk testbed to study organ preservation in a genetically human context, and that isochoric supercooling may provide a robust method for preserving and reviving engineered tissues themselves.

---

Modern organ transplantation efforts in the U.S. and abroad are hamstrung by the unavailability of on-demand donor organs^3–5^. While this may be interpreted in part as a matter of policy, the central scientific driver of this organ deficit is the extremely short *ex-vivo* shelf life of major organs. In the U.S. alone, though more than 100,000 people are currently on transplant waiting lists, more than 70% of all thoracic donor organs are discarded each year, simply due to an inability to preserve them for sufficient time periods^6^. The preservable *ex vivo* lifetime of donor hearts for instance, which are typically held on ice in “static cold storage”, is only 4-6 hours^3^.

While historical research on the preservation of organs and complex tissues focused largely on deep cryogenic temperatures, the last decade has seen the advent of “high-subzero” preservation efforts, targeting the −20° to 0°C range^7^. Chief amongst high-subzero efforts stands supercooled storage, which aims, via a number of different protocols, to hold biologics in an ice-free, metastable liquid preservation solution at sub-zero temperatures^8–10^.

In this work, we employ a recently developed technique, isochoric supercooling^2^, to achieve high-stability and predictable supercooling of the University of Wisconsin (UW) organ preservation solution at −3°C, in which we successfully preserve a functional 3D hiPSC-derived cardiac microphysiological system (MPS) for multiple days without the addition of non-physiological cryoprotectants such as dimethyl sulfoxide (DMSO) or glycerol.

The cardiac MPS combines human cells with microfluidics (Fig 1a,b) to promote 3D self-assembly into a microtissue (Fig 1c) that faithfully recapitulates complex human heart muscle structure and function^1,11–14^. The cardiac microtissues used in this work express a genetically encoded fluorescent calcium channel reporter^1^, a convenient means to optically and non-invasively track beating activity via calcium flux (Fig 1d). The calcium transients can be analyzed to obtain important information on beating activity such as beat rate, beat duration (in this study we use calcium peak duration as a proxy for action potential duration (APD)^11^) and beat shape. The MPS also allows for external electrical tissue stimulation via pacing electrodes (Fig 1d).

**Figure 1.**
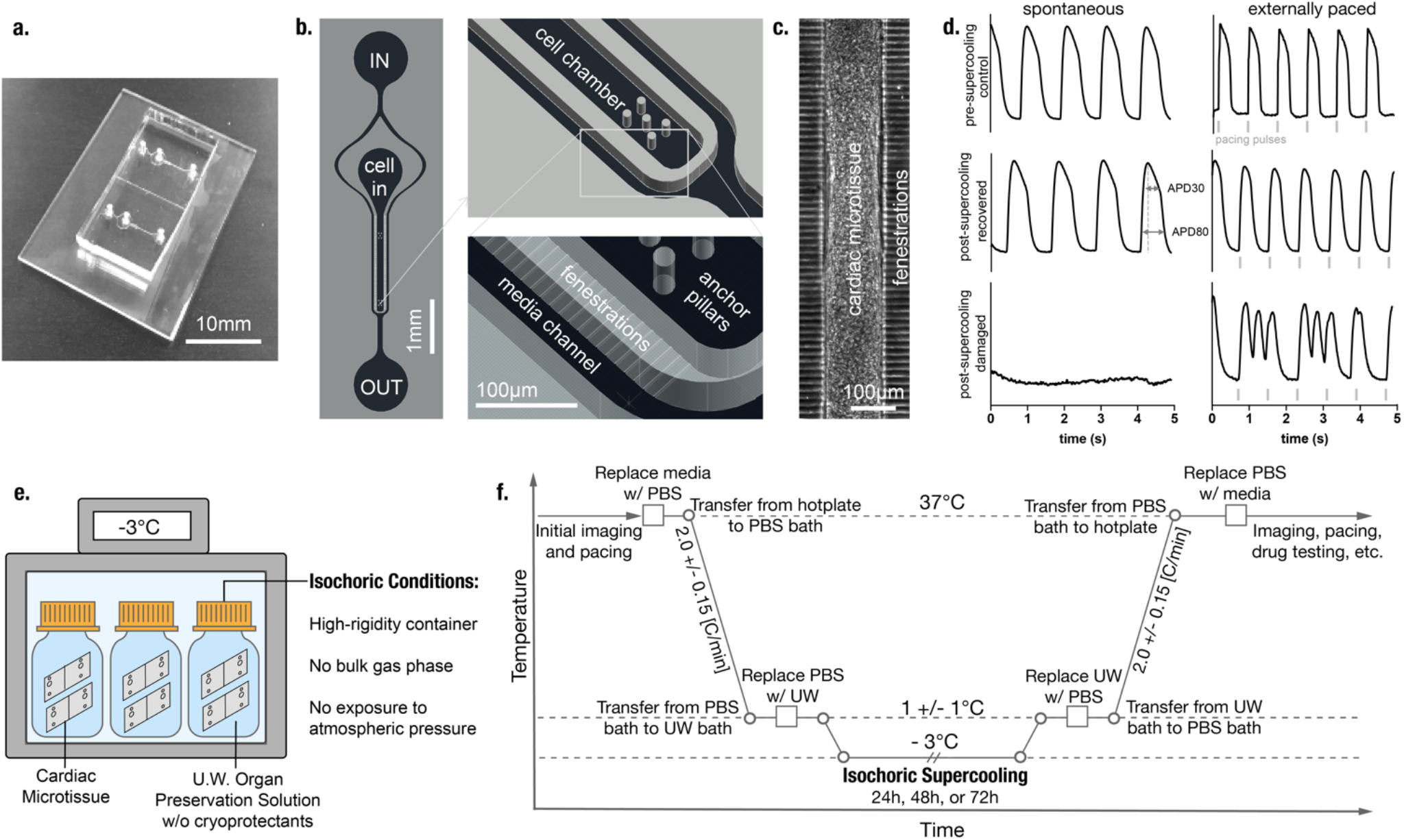
Cardiac microphysiological system and isochoric supercooling protocol. **a.** A glass-PDMS microfluidic device featuring two independent MPS. **b.** Details of the microfluidic features including media inlet (IN) and outlet (OUT) as well as the cell loading port (cell in), cell chamber with anchor pillars, media channels and connecting fenestration channels. **c.** Cardiac microtissue after cellular self–assembly within the cell chamber. **d**. Example fluorescence traces of a genetically encoded calcium channel reporter (GCaMP) showing spontaneous and externally triggered (1.25Hz) beating activity before cryopreservation as well as disturbed and restored tissue function after cryopreservation. Width of the calcium transient at 30% and 80% peak amplitude was measured as a proxy for action potential duration (APD30 and APD80, respectively). **e.** Isochoric supercooling enables preservation of biological matter in a metastable ice-free condition at temperatures below the freezing point of water / physiological saline. Isochoric conditions are achieved by confining the preservation solution in a high-rigidity container totally absent of bulk gas phase and denying it access to the atmospheric pressure reservoir, which alters both equilibrium thermodynamics and ice nucleation kinetics of the system^23,24^. **f.** Temperature-time schematic of the isochoric supercooling preservation protocol.

In designing the proof-of-concept preservation effort described herein, we established the following criteria as indicators of successful preservation: retention of sarcomere and bulk structural integrity; retention of physiological function including autonomous beating activity and responsiveness to external electrical pacing stimuli; and retention of responsiveness to a pharmacological inotrope (here the ß-adrenergic agonist, isoproterenol). Not only are these metrics rapidly monitored in the MPS, they can also be easily adapted to assess preservation success of native heart tissue or full organs in a clinical setting (e.g. QT measurements).

Inspired by previous supercooled preservation efforts involving the liver^8,9^, we designed a temperature-time trajectory by which to shuttle the cardiac MPS into and out of supercooled preservation. This trajectory is shown in Fig. 1f, and includes the following essential steps: Replacing all metabolizable media in the MPS with non-metabolizable saline solution (PBS, no calcium); submerging the MPS in a bath of PBS initially at 37°C which is then cooled to 1±1°C at a rate of approximately 2°C/min; replacing the PBS in the MPS with UW organ preservation solution (pre-chilled to 1±1°C); placing the MPS in a rigid isochoric container filled with pre-chilled UW; sealing the container tightly without trapping any bulk gas phase^2^ (Fig. 1e); and finally submerging the isochoric chamber in a constant-temperature circulating cooling bath at −3°C for the target preservation duration (24, 48, or 72 hours). Per Fig. 1f, these steps are performed in reverse for rewarming after preservation. After the MPS are returned to 37°C and their supply of metabolizable media is replenished, they are imaged once without pacing, then moved to a cell culture incubator and allowed a minimum of 24 hours of recovery before further imaging, pacing, and application of isoproterenol. A more detailed accounting of the protocol is available in the Supplementary Methods section.

The results of this preservation process are presented in Fig. 2. An average of 50-65% of tissues resumed normal spontaneous beating activity after isochoric supercooling, and a similar percentage (50-70%) responded normally to external electrical stimulation (Fig. 2a), with no significant differences observed between preservation groups. An additional 5-15% of MPS yielded partial recovery of the tissue in both cases (grey bars in Fig. 2a). Immunofluorescence staining confirmed structural integrity of the microtissue after recovery from supercooling as shown by comparable size and distribution of nuclei, as well as alignment of cardiomyocytes and sarcomeric integrity (Fig 2b). Further evaluation of physiological properties revealed that neither beat rate nor beat shape (as measured by a triangulation metric (APD80-APD30)/APD80^15^) were significantly altered by isochoric supercooling (Fig 2c,e). Beat duration (APD80) showed a slight increasing trend with increasing supercooling both in spontaneous and externally paced conditions, though any potential biological relevance of this trend (Fig 2d) is not apparent and should be explored in more detail in future studies.

**Figure 2.**
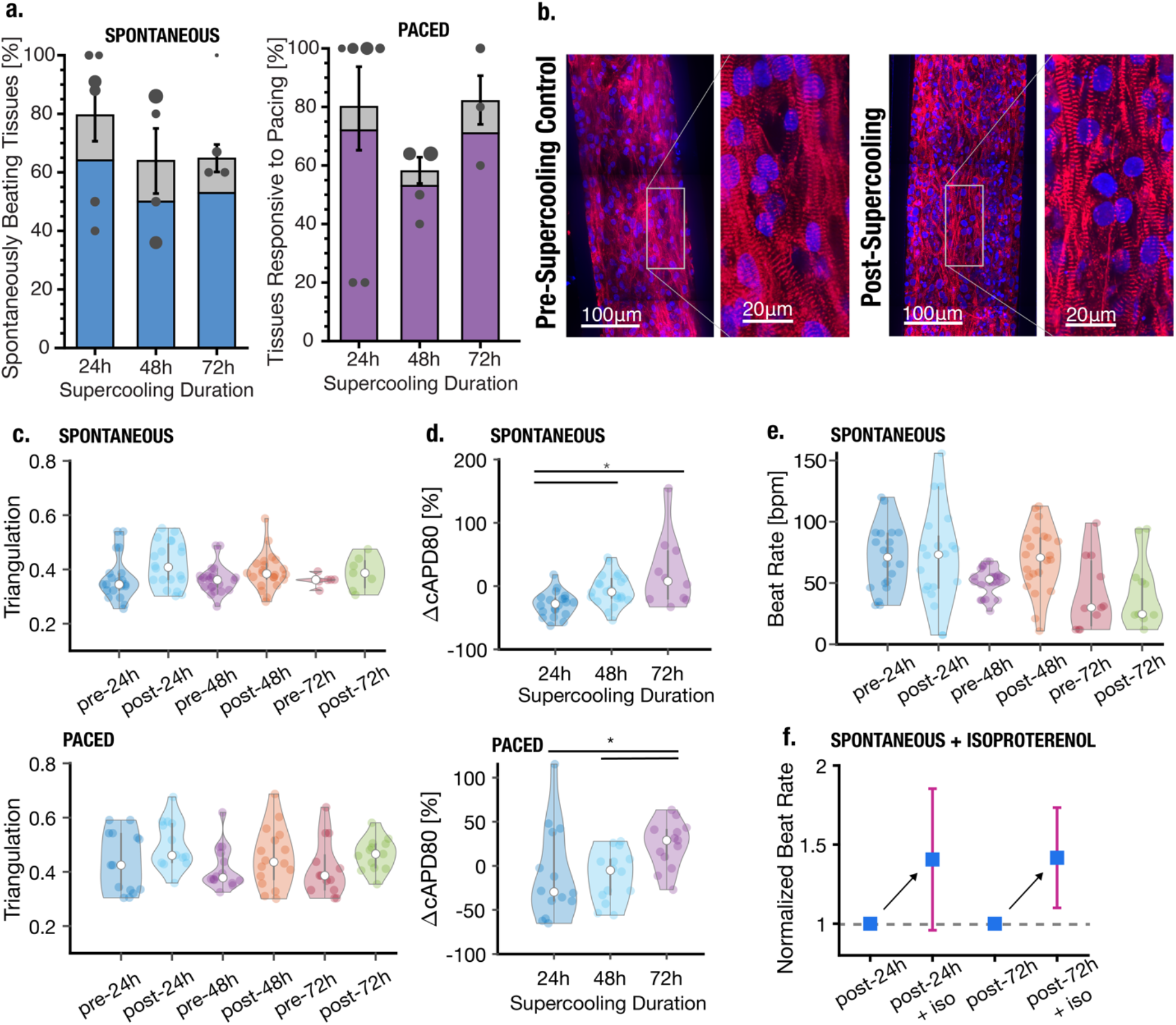
Recovery rate and comparison of key cardiac functionality parameters pre- and post-isochoric supercooling. Statistical difference between groups is marked by horizontal lines with asterisks. **a.** Total recovery rate of preserved tissues as a function of preservation time. Grey bars indicate tissues that presented some activity post-isochoric supercooling but were not coherent over the entire tissue (i.e. only part of the microtissue recovered). Scatter markers represent the weight means of individual experiments, and marker diameter reflects the number of tissues (ranging from n = 4 to n = 14 tissues). Error bars mark the standard error of the mean across experimental groups, weighted by the total number of tissues evaluated in each group. **b.** Confocal microscopy image of cardiac microtissue before and after 24h supercooled preservation (a-actinin in red and nuclei in blue). Triangulation ((APD80-APD30)/APD80) as a metric of beat shape following 24, 48 or 72h isochoric supercooling. **d.** Percent change in beat-rate corrected APD80 of recovered tissues compared to pre-isochoric supercooling. **e.** Spontaneous beat rate of recovered tissues. **f.** Increase in normalized beat rate upon introduction of 1μM isoproterenol for 30min to tissues that had previously been subjected to 24h (left) or 72h (right) supercooled preservation.

In order to probe conservation of drug response characteristics, we subjected the post-preservation MPS to 1μM of the well-known β-adrenoreceptor agonist isoproterenol. The observed increase in spontaneous beat rate further confirmed recovery of essential cardiac tissue function after isochoric supercooling, though the average beat rate increased by approximately 40% (Fig 2f), which is less than previously reported for MPS created from the same hiPSC line^12^.

With the exception of the slight increasing trend in APD80, all recovery rates, electrophysiological parameters, and drug response characteristics did not vary significantly with supercooling duration (24, 48, or 72h), suggesting that the mechanisms of damage at play are not sharply dependent on metabolism and that significantly longer preservation periods may be achievable with similarly high recovery. We hypothesize that the majority of tissue damage occurs during the cooling/warming processes, which are independent of supercooling duration and were not fully optimized in this study. Excessive cooling rates may produce damaging stresses in the tissue due to mismatch between the thermal contractility of the tissue and the surrounding microchamber. Damage may also be driven by failure to sufficiently remove metabolizable or ionic constituents prior to cooling, which can generate toxicity during rewarming due to ionic imbalances or accumulation of metabolic byproducts^16^.

Importantly, we observe no significant morphological or electrophysiological changes in recovered tissues that would suggest corruption of core cardiac functionality, as embodied most essentially by the resumption of spontaneous autonomous beating with an unchanged beat shape and expected responsiveness to isoproterenol treatment. Our data demonstrates that functional 3D microtissues may be effectively preserved at sub-zero temperatures, which has several important implications:

Firstly, these microtissue constructs may offer essential translatable insights into expected physiological effects of low-temperature preservation at the whole-organ scale. Indeed, in the drug response and pharmacokinetic analysis domains, various human-derived MPS have been demonstrated to be superior predictors of clinical response compared to animal models, offering closer physiologic and pharmacokinetic properties to native tissue whilst enabling higher-throughput testing^1,17–19^. We suggest that these same advantages can apply to low-temperature preservation, and that MPS may provide the missing link that enables timely translation of benchtop preservation protocols to the clinic, where advances in organ preservation are urgently needed.

Secondly, the high-recovery preservation results obtained using the protocol described herein, which does *not* include the introduction of non-physiological cryoprotectants, suggest that MPS platforms may also be used to study the explicit effects of various cryoprotectants on preservation efficacy. MPS platforms could be used to simultaneously screen cryoprotectants for toxicity (in a complex genetically human context), cryoprotective action, and diffusion/perfusion performance. The use of non-physiological cryoprotectants has been omnipresent in cellular preservation protocols since the 1950s, but has failed to penetrate clinical full-organ preservation, largely due to concerns surrounding toxicity or perfusion in native human tissue, and poor scaling of cell preservation results^7,20^. Our data establishes a baseline from which MPS may be used for higher-throughput, higher-reliability cryoprotectant screening.

This work also presents the first demonstration of biological preservation by isochoric supercooling^2^. This technique provides a thermodynamically simple and procedurally streamlined method of ice-free preservation at sub-0°C temperatures, and given its non-reliance on specific cryoprotective agents, we suggest that it may be applied for preservation of arbitrary biologics in the high-subzero temperature range.

In summary, this study demonstrates successful isochoric supercooled preservation and recovery of microfluidic human cardiac tissue cultures based on structural and functional characteristics. Further molecular biological studies will be needed to assess any changes on gene and biomarker protein expression levels, and optimization of thermal cycling protocols should be explored to maximize recovery rate. Moving forward, as organ-on-a-chip platforms continue to advance in both fidelity and accessibility^21,22^, we suggest that all full-organ or tissue preservation efforts should incorporate MPS preservation studies in order to probe the structural and functional effects of the preservation process with high-content, complex and human-derived tissues.

## Acknowledgements

Funding is gratefully acknowledged from the NSF Engineering Research Center for Advanced Technologies for Preservation of Biological Systems (ATP-Bio) NSF EEC #1941543. The authors thank Dr. Henrik Finsberg and Dr. Sam Wall (Simula Research Laboratory, Oslo, Norway) for the Python scripts used to extract beat information from calcium traces, Dr. Bruce Conklin (Gladstone Institutes, San Francisco, USA) for providing the hiPSCs and technical advice on the cell line and Dr. Chenang Lyu for technical assistance with the isochoric chamber assembly process.

## Author Contributions

M.J.PP., B.R., and K.E.H. conceived the study. M.J.P.P., V.C., B.C., and B.A.S. designed the preservation protocol, conducted all experiments, collected and analyzed all data, and wrote the manuscript. B.R. and K.E.H. reviewed the data and edited the manuscript.

## Data Availability

All data available upon reasonable request.

