## Supplementary Methods for "Isochoric supercooled preservation and revival of human cardiac microtissues"

### Supplementary Information

#### Methods

##### Preservation - Cooling

The cooling protocol follows the trajectory plotted in Figure 1.f. First, on a 37°C hotplate (Tokai Hit, Gendoji-cho, Japan), each MPS was flushed by gravity perfusion with 200µL phosphate buffered saline (no calcium) in order to remove metabolizable compounds from the MPS prior to cooling. This step was performed to minimize generation of metabolic byproducts during cooling that may accumulate during storage and cause toxicity upon return to normothermia. Importantly, PBS without calcium was employed during cooling and warming in order to minimize the risk of calcium shock or other ionic imbalance effects as tissues warm up and get reperfused with culture media. The surface of the MPS device was briefly wiped with 70% Ethanol to reduce contamination risks. The MPS were then submerged in a well of approximately 30mL of PBS at 37°C. This well was then placed directly into a circulating ice and water bath at 1±1°C, resulting in a cooling rate of 2.1±0.15°C per minute. The temperature in both the well and a reference MPS were continuously monitored via thermocouple. When the reference MPS temperature reached below 2°C, all MPS were submerged in an identical well filled with equally cold University of Wisconsin (UW) organ preservation solution. The MPS were then flushed with 200µL of pre-chilled UW solution by gravity perfusion.

##### Preservation – Storage under isochoric supercooling

Following cooling, the MPS were transferred to an approximately 65 mL thick-walled glass container filled with pre-chilled UW solution at 1±1°C. Crucially, care was taken to ensure that no bulk gas phase (e.g. air bubbles) were present in the solution before and after introduction of the MPS. The container was filled to the brim and then sealed using a rigid threaded polypropylene cap that included a centered plug to displace air and liquid as the cap is threaded down, and ensures that the sealed container contains no air. A more detailed accounting of this assembly process, the described components, and the thermodynamic and kinetic effects of confining the preservation liquid in constant-volume (isochoric) conditions absent air are available in our previous publication[1].

The isochoric container, housing 2 – 8 MPS per container, was then submerged in a constant-temperature circulating cooling bath (PolyScience, USA) at -3°C for 24, 48, or 72 hours. This preservation temperature was chosen based on our previous isochoric nucleation experiments showing that isochoric supercooling was extremely stable (e.g. the likelihood of ice nucleation was <1%) at -3C for physiological saline, which has a near-identical freezing point to UW solution (-0.56°C). The stability of isochoric supercooling is a multifaceted consequence of denying the system access to a pressure reservoir (e.g. the atmosphere), which suppresses density fluctuations[1]; denying the system access to air-water interfaces, which may function as nucleation sites[1,2]; and suppressing secondary nucleation mechanisms such as cavitation-induced nucleation[3].

##### Preservation – Warming

Per Fig. 1.f., in order to warm the MPS back to physiological temperatures, the cooling protocol was applied precisely in reverse. It should be noted in particular that the isochoric container was warmed from -3°C back to 1±1°C (via ice bath) before it was opened. Premature opening of the isochoric container (and disruption of isochoric thermodynamic conditions) at temperatures less than -0.56°C can lead to immediate ice formation.

##### Recovery

In initial protocols, the MPS were imaged for spontaneous activity on a hotplate directly following warming and before being placed back into an incubator at 37°C with (5% $CO_2$). Of note, in this directly post-warming measurement, incoherent electrophysiological activity was visually observed in several MPS. Constellations of calcium transients could be observed in various parts of the tissue, even while coherent signaling across the whole tissue (as accompanies autonomous beating) was absent. Early in our study design, this observation indicated to us that tissues may not be dead, even while not spontaneously beating, and we thus elected to employ a minimum-24h recovery period before electrical pacing and further imaging. Later experiments had an adjusted protocol where MPS were directly put in the incubator after warming (skipping the previously described imaging), to rapidly expose the tissues to physiological temperature and $CO_2$. The results reported in Fig. 2 all reflect incubated recovery times of at least 24h (minimum 24h, maximum 96h). We did not distinguish between recovery times in aggregating the data presented in Fig. 2, as statistical analysis of the recovery rate based on supercooling duration and recovery time showed no significant difference between recovery times greater than 24h. However, while outside of the scope of this study, dedicated high-time-granularity study of the recovery process within the first 24 hours post-supercooling may yield important insight into the physiological effects of thermal cycling and prolonged hypothermic storage and the cardiac recovery process. Similarly, continued observation beyond 96h post-preservation will give insight into the long-term survival and stability of previously supercooled MPS.

#### Microfluidic device design and fabrication

Cardiac MPS were designed and prepared as previously described[4–6]. In brief, the MPS were fabricated using traditional soft lithography. Polydimethylsiloxane (PDMS; Dow Chemical, Sylgard 184) was prepared as 10:1 mixture with curing agent and poured onto silicon wafers that served as master molds featuring the negatives of the microfluidic structures made from SU-8. After curing, the PDMS was peeled off the master mold and cut to size. Holes of 0.75 mm diameter were punched out for access ports. Oxygen plasma surface treatment (21W, 24s, 600mTorr) was applied to permanently bond the PDMS to 1mm glass slides (VWR, 48300-048) cleaved to 2.5x1.5 cm (Figure 1A).
The microfluidic design featured elongated cell chambers that promote cellular self-assembly into a uniaxially beating microtissue (Figure 1B). The chamber dimensions varied from 800 to 1800µm length; 120 to 170µm width and 60 to 70µm height. Media channels ran in parallel on either side of the cell chamber. Media exchange in the cell chamber was achieved via an array of small connection channels (2x2µm cross-section; 40µm length) that protected the tissue from fluid mechanical forces[4]. Small pillars (20µm diameter) at either end of the cell chamber served as structural anchor points to keep the muscle fiber from collapsing. Each microfluidic device featured either a single tissue chamber or 4 identical tissue chambers in parallel.

#### CM differentiation from iPSC

Cell culture was performed similar to already published protocols[6]. We used the human iPSC line Wild Type C (WTC; # GM25256, Coriell Institute) edited to express a fluorescent reporter for intracellular calcium (GCaMP6f)[7]. After thawing the iPSCs were cultured on Matrigel (Corning, 354277) in mTeSR-1 media (Stem Cell Technologies, 85851) for several passages every three days before differentiation was started. Accutase (Millipore, SCR005) was used for dissociation. Differentiation was induced via the Wnt/ß-catenin signaling pathway. First, 8µM CHIR 99021 (Peprotech, 2520691-10MG) was applied for 24h in RPMI 1640 media (Gibco, 11875-093) with 2% of B-27 Minus Insulin (B27-I, Gibco, A18956-01), followed by 24h in RPMI + B27-I without any small molecules. Then, 5µM IWP-4 (Peprotech, 6861787-10MG) was applied for 48 hours, followed by 48h in RPMI + B27-I. CM differentiation was completed with subsequent 48 hour media exchanges with RPMI + 2% B27 supplement that contained insulin (B27+C, Gibco, 17504-044) until the cells began to beat spontaneously. CM were used up to 14 days after differentiation.

#### Cardiac MPS loading and maintenance

For MPS loading, cells were singularized using collagenase type 2 (Worthington Biochemical Corporation, LS004176) for 45-60 minutes in a cell culture incubator and collected in EB20 media (Knock Out DMEM (Gibco, 10829-018) with 20% FBS (Gibco, 16000-044), 1% MEM non-essential amino acids (Gibco, 11140-050), 1% Glutamax (Gibco, 35050-061) and 400nM 2-Mercaptoethanol (Gibco, 21985-023)) supplemented with 10µM ROCK-inhibitor (RI, Y-27632 dihydrochloride, Peprotech, 1293823-10MG). Each cell chamber was loaded with 3µl cell suspension containing enough cells to completely fill the chamber (between 5000 and 20000 cells). The cell suspension was injected into the cell loading port using a pipette tip (Rainin, LTS ultra fine 20µL), which was cut to about 1cm length and remained in the chip. Two centrifugation steps (3min at 300 rcf; first horizontal then vertical orientation of the chip) were performed to aggregate the cells and move them into the cell chamber. The cell loading port was then sealed and EB20 + RI media was perfused through the media channels after 1h incubation time. The next day media was exchanged to RPMI + 2% B27+C and changed 3 times per week thereafter. MPS were used within 30 days after loading.

#### Data acquisition

Videos were obtained using a digital CMOS camera (HAMAMATSU, C11440 / ORCA-Flash 4.0) mounted to Nikon TE-300 inverted microscope with an LED light engine (Lumencor SpectraX). A heated platform (Tokai Hit, TPi-SQX) was used to image the MPS at 37°C. For electrical stimulation a pulse generator (ION OPTIX Myopacer Field Simulator) was used. 20V biphasic rectangular pulses (20ms) were applied via 1.5" long blunt stainless-steel needles (Vita Needle M937) inserted into the media inlet and outlet pipette tips. Calcium traces were recorded at 100fps with 4x4 binning using the cyan LED for excitation. A 1mM isoproterenol stock was prepared freshly from isoproterenol hydrochloride (I0260; TCI Chemicals) powder right before the drug response experiments, sterile filtered and diluted to 1µM. 100µL of drug-containing media were applied to each chip via the inlet pipette tip 30min before recording.

#### Data analysis

Fluorescence videos were converted into time-intensity profiles (Figure 1D) using an in-house library of python scripts. Beat parameters such as beat rate and peak duration at 30% and 80% intensity (APD30 and APD80, respectively) were automatically detected. Further metrics were calculated: The beat-rate corrected peak duration (cAPD80) was computed using Fridericia method, which corrects for variations in beat rate by scaling the values to 1Hz beat rate[8]: $cAPD_{80} = APD_{80}/BR^{-\frac{1}{3}}$ with BR being the beat rate in Hz and $cAPD_{80}$ being the peak duration at 80% peak height corrected to a beat rate of 1Hz. Triangulation, as introduced by Hondeghem et al.[9], was computed as a metric of beat shape: $(APD_{80} - APD_{30})/APD_{80}$. Lower triangulation values indicate a more rectangular beat shape while higher values indicate stronger triangulation, which is associated with increased pro-arrhythmic potential.

For analysis of the recovery rate we counted the percentage of tissues that resumed beating activity after preservation as judged by contractile tissue motion or peaks in the calcium trace. We distinguished between samples that showed coherent beating of the entire tissue as one unit from those that showed activity only in a portion of the tissue. We defined recovery rate as the number of tissues that resumed beating activity spontaneously or in response to electrical stimulation at any time point after preservation, divided by the total number of preserved tissues.

Graphs were plotted in MATLAB (version R2019b). Figures were assembled using Adobe Illustrator (version 25.0.1). Statistical analysis to compare preservation effects was done by one-way ANOVA, with Bonferroni's method used for multiple comparisons test (all performed in MATLAB using built-in functionality). Post-preservation results were compared to pre-preservation results for each condition, as well as to other post-preservation results at different supercooling timepoints. Effects were reported for the significance levels of $p < 0.05$ (*). In the presented data, horizontal bars topped by an asterisk indicate significant difference between groups, and if no bars are shown, the groups were determined to be statistically similar.

#### Immunofluorescence

After preservation the MPS were allowed to warm up and resume beating. After 24h of recovery, tissues were washed with phosphate buffered saline (PBS) for 5min and then perfused with 4% paraformaldehyde (PFA) for 15min. After another two washing steps with PBS, the PDMS was carefully filleted off the glass slides using a scalpel to expose the tissues. The tissues, which remained attached to the PDMS, were sequentially submerged in the following solutions: blocking buffer (BB: 1% BSA 10% FBS 0.5% Triton 0.05% sodium azide) overnight at 4°C; DAPI 1:1000 (Invitrogen D1306) in BB for 30-40min at 25°C; primary antibody (Mouse anti α-actinin, Life technologies 41811) 1:100 in BB for 48h at 4°C; two washing steps in BB for 2h at 25°C and 1 washing step over night at 4°C; secondary antibody (Goat anti-mouse IgG Alexa 568 H+L, Life Technology a11004) 1:100 in BB overnight at 4°C; two washing steps in BB for 2h at 25°C and 1 washing step over night at 4°C. All incubation steps were performed on a shaker. Tissues were imaged in the Opera Phenix™ High Content Screening System using a Proprietary Synchrony™ Optics 63x water immersion lens and Harmony acquisition software. We performed z-stacks over 60µm height with a step-size of 0.5µm. Image processing in ImageJ was done to stitch large images, perform maximum intensity projection (15 slices) and enhance contrast.
